# Surface Electromyography for Identification of Pre-Phonatory Activity

**DOI:** 10.1101/2021.07.29.454386

**Authors:** Hardik Kothare, Mark S. Courey, Katherine C. Yung, Sarah L. Schneider, Srikantan Nagarajan, John Houde

## Abstract

Surface electrode EMG is an established method for studying biomechanical activity. It has not been well studied in detecting laryngeal biomechanical activity of pre-phonatory onset. Our aims were to compare the sensitivity of surface EMG in identifying pre-phonatory laryngeal activity to needle electrode laryngeal EMG and to compare the pre-phonatory period in patients with adductor laryngeal dystonia (ADLD) with that in controls. ADLD patients (n = 10) undergoing needle LEMG prior to Botox^®^ injection and participants with normal voices (n = 6) were recruited. Surface EMG electrodes were placed over the cricoid ring and thyrohyoid membrane. Needle EMG electrodes were inserted into the thyroarytenoid muscle. EMG and auditory output samples were collected during phonation onset. Tracings were de-identified and evaluated. Measurements of time from onset in change of the amplitude and motor unit frequency on the interference pattern to onset of phonation were calculated by two blinded raters. 42 of 71 patient and 40 of 50 control tracings were available for analysis. Correlation for pre-phonatory time between electrode configuration was 0.70 for patients, 0.64 for controls and 0.79 for all the data combined. Interrater correlation was 0.97 for needle and 0.96 for surface electrodes. ADLD patients had a longer pre-phonatory time than control subjects by 169.48ms with surface electrode and 140.23ms with needle electrode (p < 0.001). Surface EMG demonstrates equal reliability as Needle EMG in detecting pre-phonatory activity in controls and subjects. Patients with ADLD have a significantly prolonged pre-phonatory period when compared with controls.

## Introduction

Surface electrode electromyography (SEMG) is a well-established technique for studying biomechanical activity. By placing surface electrodes on the skin over the belly of a muscle, the electrical activity generated during contraction can be assessed. In risk of oversimplification, the fidelity and the amount of activity detected depends on how close the electrodes are to the active region in the target muscle, the type and configuration of the surface electrodes used, the force of contraction of the muscle and potential interference (crosstalk) generated from competing muscles in the region (1).

Regarding laryngeal activity and SEMG, except for the cricothyroid muscle, the intrinsic muscles of the larynx are separated from the skin by the thyroid cartilage and cervical strap muscles. Therefore, conceptually using SEMG to study biomechanical activity of the intrinsic laryngeal muscles is potentially limited by crosstalk from the muscles between the skin and the laryngeal framework. With this understanding, most of the existing research on SEMG and voice has primarily evaluated activity of the strap muscles, tongue muscles and other cervical muscles in patients with dysphonia (2). To our knowledge SEMG has not been used to study the biomechanical activity of the intrinsic laryngeal muscles for the onset of phonation.

Prior to the onset of voice, the intrinsic laryngeal muscles must be activated to place the mucosa into a position appropriate for phonation. This biomechanical activity known as pre-phonatory activity or pre-phonatory burst activity has been demonstrated and evaluated by multiple laryngeal electromyographic studies with needle electrodes inserted directly into the intrinsic laryngeal muscles (3-5). This procedure is often painful and perceived as invasive. The invasiveness and pain from needle EMG are barriers into research on the relationship between pre-phonatory activity and voice production in patients with neurological disorders that may affect voice onset and maintenance such as in patients with laryngeal dystonia (LD).

The purpose of this study was to evaluate the specific ability of SEMG to evaluate phonation onset reliably and precisely. Our goals were: 1. to determine if EMG performed with surface electrodes was as sensitive at identification of pre-phonatory laryngeal adductor activity as EMG performed with indwelling needle electrodes; and 2. to compare the length of the pre-phonatory period between individuals with a voice disorder of neurological origin, adductor laryngeal dystonia (ADLD), to participants with normal voices.

## Methods

Informed consent was obtained from all participants for these experiments following institutional approval by the Committee for Human Research at UCSF. Ten patients with ADLD (5 males/5 females with an average age of 56.10 +/- 13.06 years) uncomplicated by the co-existence of dystonic tremor or chronic vocal strain as judged by perceptual analysis were recruited from the UCSF Voice Clinic. All patients had exhibited repeated prior good response to Botox^®^ injection. At the time of collection of EMG data, all patients were experiencing a return of ADLD symptoms. They had presented to the clinic for repeat Botox^®^ injection under monopolar needle laryngeal EMG for guidance during the injection of Botox^®^. Six controls (4 males/2 females with an average age of 44.17 +/- 9.33 years) were also recruited. All participants without LD were judged to have normal voice by perceptual analysis performed by a Speech-Language Pathologist and a laryngologist each with greater than 15 years of experience.

Active and reference surface EMG electrodes (CareFusion Stainless Steel Disc Electrodes 6030-3-TP, Middeton, WI) were placed over the cricoid ring and thyrohyoid membrane. For all surface electrodes, electrode paste was applied to the skin at the site of electrode contact (Parker Laboratories, Inc, Fairfield, NJ). For direct muscle samples the active monopolar 27 gauge needle electrode (Natus Manufacturing Ltd, Galway, Ireland) was inserted directly into the thyroarytenoid muscle, while the reference surface electrode (CareFusion Stainless Steel Disc Electrodes 6030-3-TP, Middeton, WI) was placed over the clavicular head. For both sets of electrodes, a single surface ground electrode was placed over the patient’s forehead. The 2 electrode configurations, along with the audio signal, were hooked to a 4-channeled bridge (Nicolet Biomedical, EA-4, 1404262, Madison, WI) connected to a Nicolet Viking EMG unit (Nicolet VikingSelect G2, NK030199R. Madison, WI). The signals were linked in time, filtered, and displayed at a sweep speed of 100 milliseconds. For surface electrodes, the display amplitude was constant at 100 microvolts per division. For the needle electrode configuration, the display amplitude was varied between 200 and 500 microvolts per division to prevent contamination by other signals (Figure 1).

**Figure 1.**
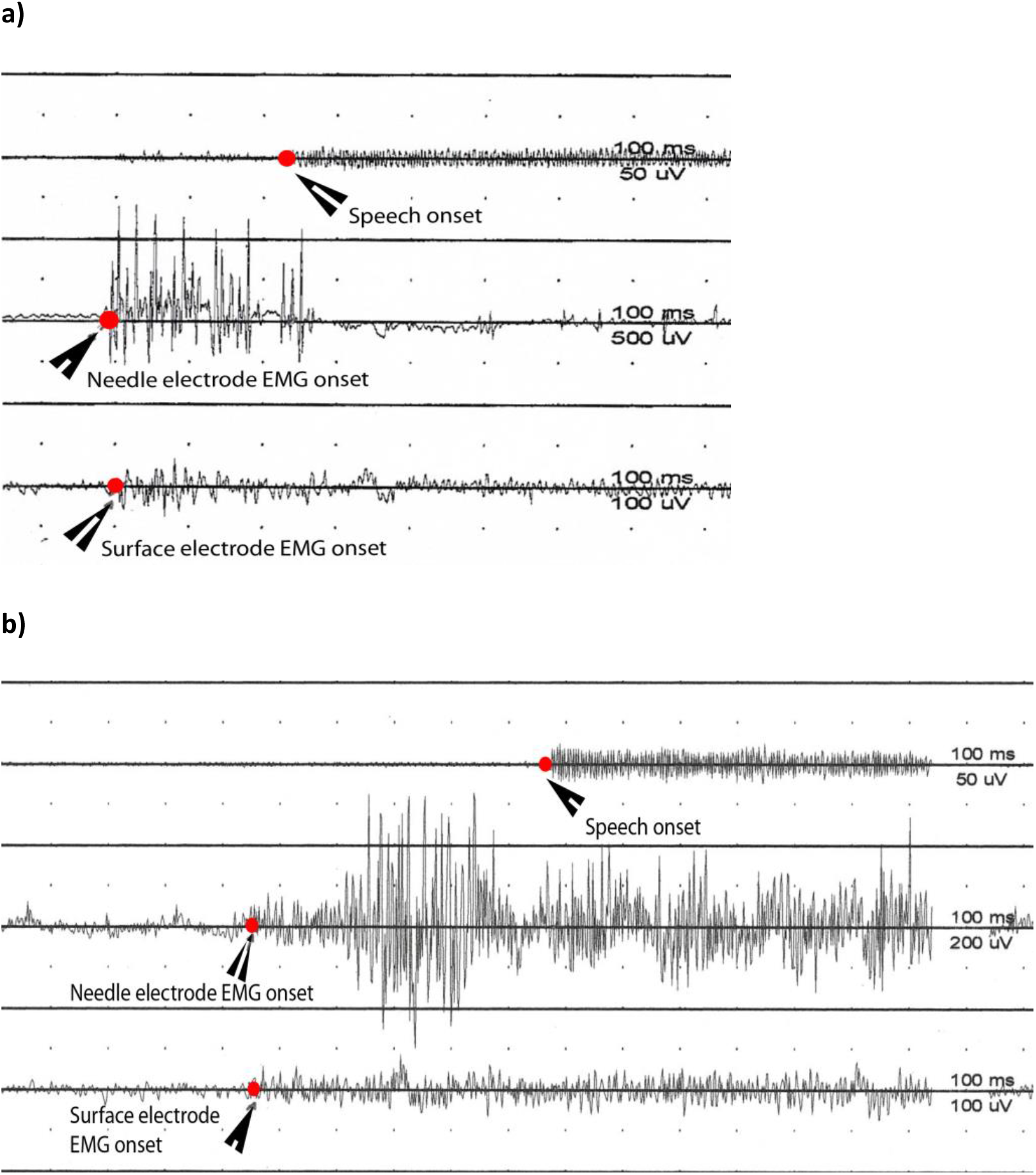
**a)** Paired tracing from a control. Top line is the phonatory signal. Middle line is the needle EMG signal. Bottom line is the surface EMG signal. **b)** Paired tracing from a patient with ADLD. Top line is the phonatory signal. Middle line is the needle EMG signal. Bottom line is the surface EMG signal. The elongated pre-phonatory interval can be seen in this patient both in the SEMG and needle EMG measurements.

With the electrodes in place, participants were asked to phonate /i/ at a comfortable pitch and loudness level. Computer screens of the interference patterns were captured during the period of phonation onset and later printed for analysis. Attempts were made to capture baseline activity followed by onset of activation. 3 to 5 samples were collected for each experimental subject and control. These samples were collected in one session on one day lasting 5 to 7 minutes.

Printed EMG and auditory output samples were de-identified and split so that each tracing contained the auditory output signal and either the needle electrode interference pattern or the surface electrode pattern. These tracings were evaluated visually by 2 blinded reviewers to detect change in amplitude and/or motor unit density of the EMG signal. Using the time scales on the tracings, the time in milliseconds from the onset in change of the amplitude and motor unit frequency on the interference pattern to the onset of phonation were taken from each recording. Differences in ratings from each sample for each of the 2 reviewers were averaged and the variance calculated to determine interrater reliability. Acceptable variance in pre-phonatory time measures between raters was set at 10%. Linear regression analysis was used to calculate Pearson’s correlation coefficient to determine linear association between surface electrode onset and needle electrode onset. An unpaired t-test assuming unequal variance (α = 0.05) was used to compare the length of the pre-phonatory period between patients and controls.

## Results

### Tracings

Attempts were made to capture 3 to 5 tracings per subject and control. From the 10 patients, there were 71 total tracings. Of these 71, 39 were adequate for evaluation in the surface electrode configuration and 37 in the needle electrode configuration. From the 6 controls, there were 50 total tracings. 35 were adequate for analysis from the surface electrode configuration and 22 from the needle configuration. Tracings were excluded from analysis due to low signal-to-noise-ratio (patients 14, controls 6), excessive gain on needle electrode channel that caused overlapping of tracings (patients 12, controls 0), phonation onset was not captured in the tracing (patients 1, controls 2), pre-phonatory motion artifact created an unstable baseline (patients 8, controls 2), and on some tracings the pre-phonatory burst could not be identified (patients 1, controls 5). The differences in the number of tracings available for analysis was not statistically different between patients and controls (Fisher’s exact test, p=0.130 for surface electrode configuration and p = 0.461 for needle electrode configuration).

### Inter rater reliability

Raters visually examined the signals and determined the time of onset of EMG activity and the voice signal. They each then determined the length of the pre-phonatory time period using the time scale from the tracing. Allowing 10% overall variability, inter rater correlation was 97% for the needle electrode configuration and 96% for the surface electrode configuration.

### Pre-phonatory times can be accurately measured with surface EMG

Linear regression analysis for pre-phonatory time between electrode configurations revealed an R-squared value of 0.70 for patients, 0.64 for controls and 0.79 for all the data combined. (Figure 2).

**Figure 2.**
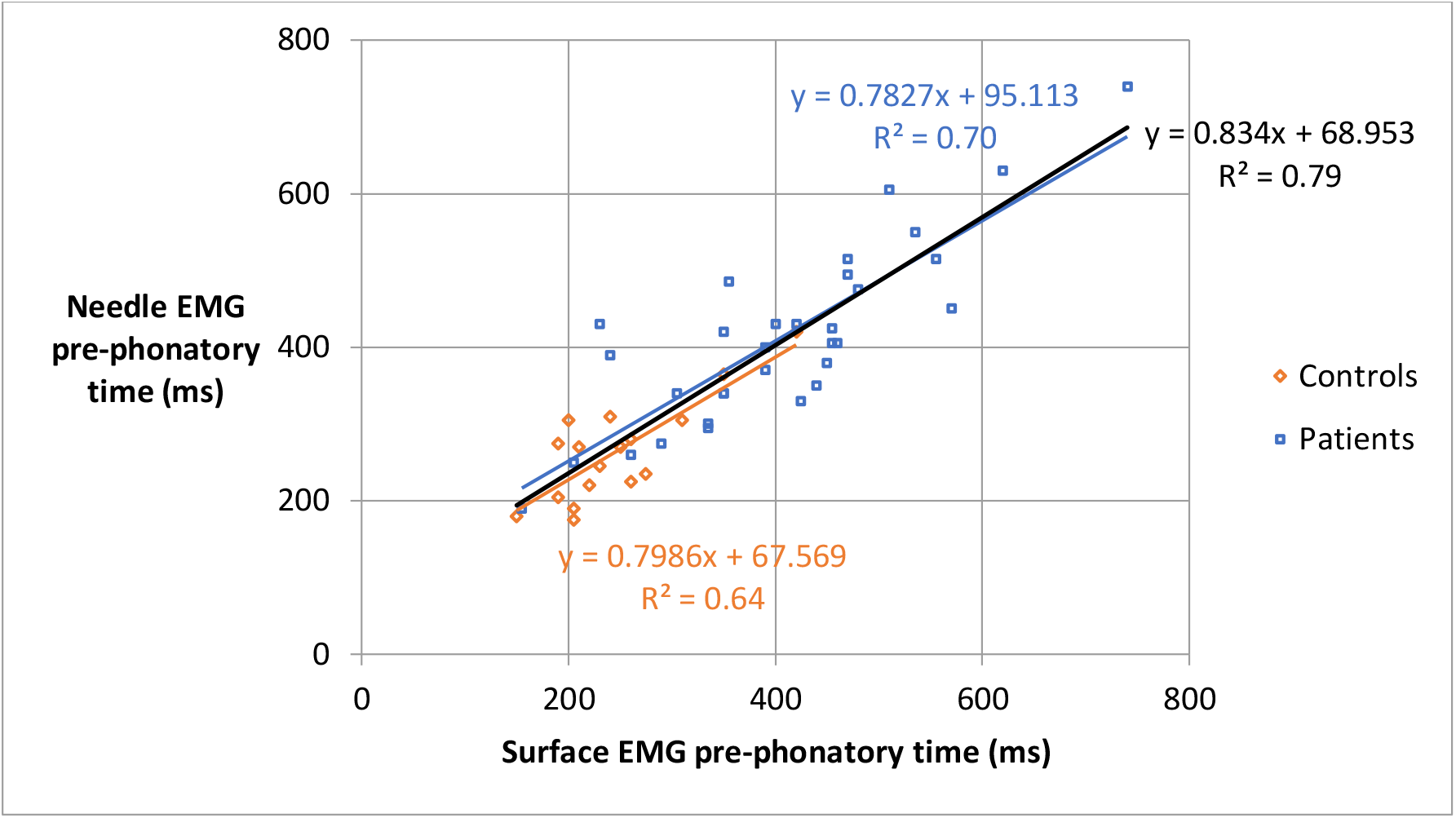
Scatter plot of Linear Regression Analysis for pre-phonatory time between electrode configurations

### Both Needle and Surface EMG show elongated pre-phonatory times in LD

The average pre-phonatory time for the control group was 245 +/- 9.5ms in the surface electrode configuration and 271 +/- 14ms in the needle electrode configuration. This was not significantly different (unpaired t-test, p=0.132). In patients, the average pre-phonatory time was 414 +/- 20ms in the surface electrode configuration and 411 +/- 19ms in the needle electrode configuration. This was not significantly different (unpaired t-test, p=0.908). Patients with adductor LD had a longer pre-phonatory time than control patients by 169.48ms with surface electrode and 140.23ms with needle electrode which was significantly different in both electrode configurations (unpaired t-test, p < 0.001) (Figure 3).

**Figure 3.**
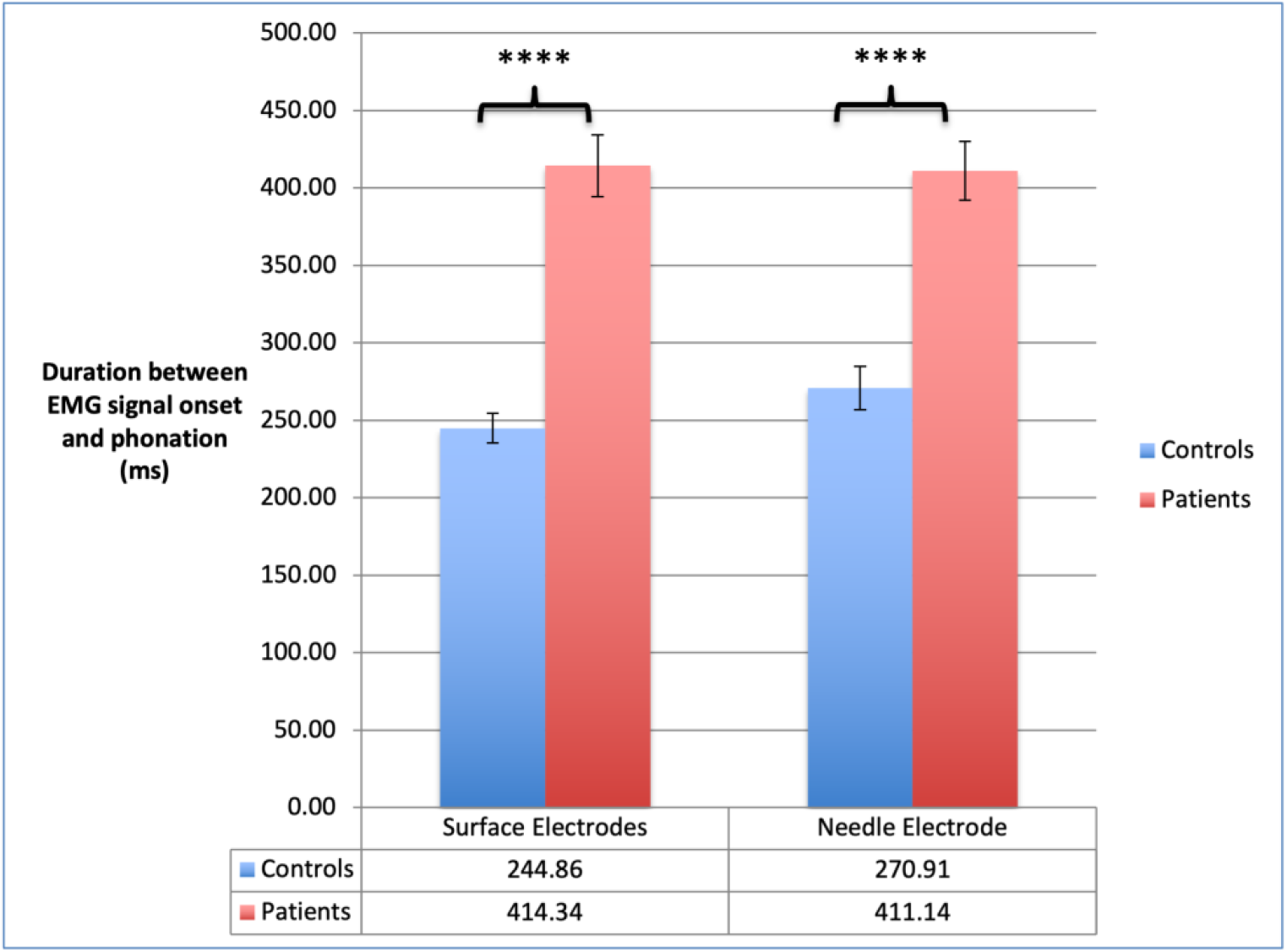
Comparison of pre-phonatory time between electrode configurations and participants.

## Discussion

SEMG is an established method for the study and evaluation of biomechanical activity. It is most commonly used to study activity in muscles located just below the skin. Within the head and neck, SEMG has been used primarily to study strap muscle (infrahyoid and suprahyoid) activity related to swallowing and speech production in hypofunctional voice disorders. To our knowledge, SEMG has not been used to study intrinsic laryngeal activity. Due to the distance of the intrinsic laryngeal muscles from the surface of the skin, and the interposition of muscles between the skin and larynx, it is not intuitively obvious that SEMG would be adequate to assess intrinsic laryngeal biomechanical activity. Prior research has shown that pre-phonatory activity can be assessed accurately with needle electrodes. However, needle electrodes are invasive and painful. This is a barrier to research into the pre-phonatory period. Our goal was to develop a less invasive method to study this activity. As such we compared surface electrode activity with needle electrode activity. Our results show that the ability of these different electrode configurations to detect the pre-phonatory activity is comparable. With our technique, we were able to capture pre-phonatory activity with similar reliability with either SEMG or Needle EMG. Furthermore, when identified, there was excellent agreement between SEMG and Needle EMG configurations. It is likely that the EMG electrodes picked up crosstalk from activity in the strap muscles. However, the strong correlation with activity detected by Needle EMG suggests that surface EMG signals correlate with intrinsic laryngeal activity. This finding helps establish the validity of SEMG to identify and study the pre-phonatory period.

The validity of our findings is further supported by the ability of both electrode configurations to identify prolongation of the pre-phonatory activity in patients with adductor laryngeal dystonia as compared to participants without voice disorders. This abnormality has been identified previously by other researchers (6). As our findings are consistent with theirs, it is likely that the activity we are seeing is a correlate of pre-phonatory activity within the intrinsic laryngeal muscles, if not laryngeal adduction itself. Taken together these findings support the validity of SEMG for accurately identifying the intrinsic laryngeal muscle biomechanical activity that occurs during the pre-phonatory period.

## Conclusions

SEMG demonstrates equal reliability with Needle LEMG in detecting laryngeal pre-phonatory activity in controls and patients. Patients with ADLD have a significantly prolonged pre-phonatory period when compared with aged-match controls.

